# Rapid and stable microbial community assembly in the headwaters of third-order stream

**DOI:** 10.1101/426460

**Authors:** Morgan E. Teachey, Jacob M. McDonald, Elizabeth A. Ottesen

## Abstract

Small streams and their headwaters are a key source of microbial diversity in fluvial systems and serve as an entry point for bacteria from the surrounding landscape. Community assembly processes occurring in these streams shape downstream population structure and nutrient cycles. To elucidate the development and stability of microbial communities along the length of a first through third order stream, fine-scale temporal and spatial sampling regimes were employed along McNutt Creek in Athens, Georgia, USA. 16S rRNA gene libraries were constructed from samples collected on a single day from 19 sites spanning the first 16.76 km of the stream. Selected sites at the upper, mid, and lower reaches of the stream were sampled daily for 11 days to evaluate community variability over time. In a second study, sites at and near the creek’s headwaters were sampled daily for 11 days to understand the initial stages of bacterioplankton community assembly. In all studies, we observed decreasing alpha and beta diversity with increasing downstream distance. These trends were accompanied by the enrichment of a small fraction of taxa found at low abundance in the furthest-upstream environments. Similar sets of taxa consistently increased significantly in relative abundance in downstream samples over time scales ranging from 1 day to 1 year, many of which belong to microbial clades known to be abundant in freshwater environments. These results underpin the importance of headwaters as the site of rapid in-stream selection that results in the reproducible establishment of a highly stable community of freshwater riverine bacteria.

**Importance:** Headwater streams are critical introduction points of microbial diversity for larger connecting rivers and play key roles in the establishment of taxa that partake in in-stream nutrient cycling. We examined microbial community composition of a first- through third-order stream using fine-scale temporal and spatial regimes. Our results show that the bacterioplankton community develops rapidly and predictably from the headwater population with increasing total stream length. Along the length of the stream, the microbial community exhibits substantial diversity loss and enriches repeatedly for select taxa across days and years, although the relative abundances of individual taxa vary over time and space. This repeated enrichment of a stable stream community likely contributes to the stability and flexibility of downstream communities.

## Introduction

Riverine systems act as an interface between many distinct habitat types, connecting hillslope and bottomland soils to downstream bodies of water and controlling the flow of bacteria and nutrients from one environment into the next. In many fluvial systems, alpha and beta diversity both typically decrease along the length of a stream or stream network (1-11). This suggests that headwater streams are a critical source of microbial biodiversity. The primary source of microbes entering headwater bacterial communities appears to be soil and soil waters (1, 2, 8, 12), with pelagic stream community assembly resulting from the enrichment of bacteria present in these environments at low abundance. From the headwaters, the bacterioplankton community continues to develop as it travels through the stream network (10, 11, 13).

Pelagic stream communities are renewed continuously via the paired forces of in-stream selection and bulk transport of organisms from upstream and upslope environments. This raises the possibility of rapid and significant shifts in community composition within streams resulting from fluctuations in physicochemical conditions, flow rates, and the composition or origin of microbes entering the stream. Temporal trends in community profiles have been described in many riverine systems (3, 5-7, 12-18). In several of these studies, the same taxa were found over time, varying in abundance based on shifts in physiochemical factors or following landscape-level disturbances (5, 10, 13, 17-20). This reoccurrence of taxa has led several groups to suggest the presence of a core community that is of particular importance in shaping riverine community dynamics (2, 9, 10, 13, 21-23). Conversely, multiple studies have also found substantial variation in microbial community structure and function over time in a wide range of study systems (3, 5, 6, 10, 15-18, 20, 22, 24). A temporal study of headwater streams and an adjoining higher-order stream by Portillo et al. (24) found seasonal variation among samples taken from the same site. Despite their proximity, samples clustered by site regardless of collection day (24). These findings indicate fluvial core community may either fluctuate over time, or that changing environmental conditions can lead to alterations in the relative abundance of core vs. transient community members.

Few studies have evaluated the extent of short-term variability between the microbial communities of headwater streams, as well as how daily fluctuations in upstream community composition may impact downstream community structure and function. Further investigation into the assembly and stability of small stream bacterioplankton populations on both fine temporal (days) and spatial scales is necessary to advance the current understanding of the initial development of freshwater pelagic communities. Fundamental questions remain, including: 1) To what extent are fine-scale spatial patterns of community diversity and composition reproducible over time?; and 2) Do in-stream communities exhibit a temporal memory, or distance-decay relationship, and at what time scale(s)? To directly assess this knowledge gap, two multi-day studies on a small creek in Athens, GA, USA were performed.

Our chosen study site, McNutt Creek (Figure 1A), has consistently demonstrated a loss of diversity with increasing dendritic distance as well as a corresponding increase in common freshwater taxa across seasons, as seen in larger riverine systems and higher-order streams (25). These characteristics make McNutt Creek an ideal study system to address these questions regarding bacterial community assembly in headwater streams. In this work, a rapid rise in dominance of freshwater-associated bacterial taxa along the stream flow path was observed. Community assembly was renewed daily and quickly recovered following landscape-level disturbance, revealing robust community stability across the stream’s flowpath.

**Figure 1:**
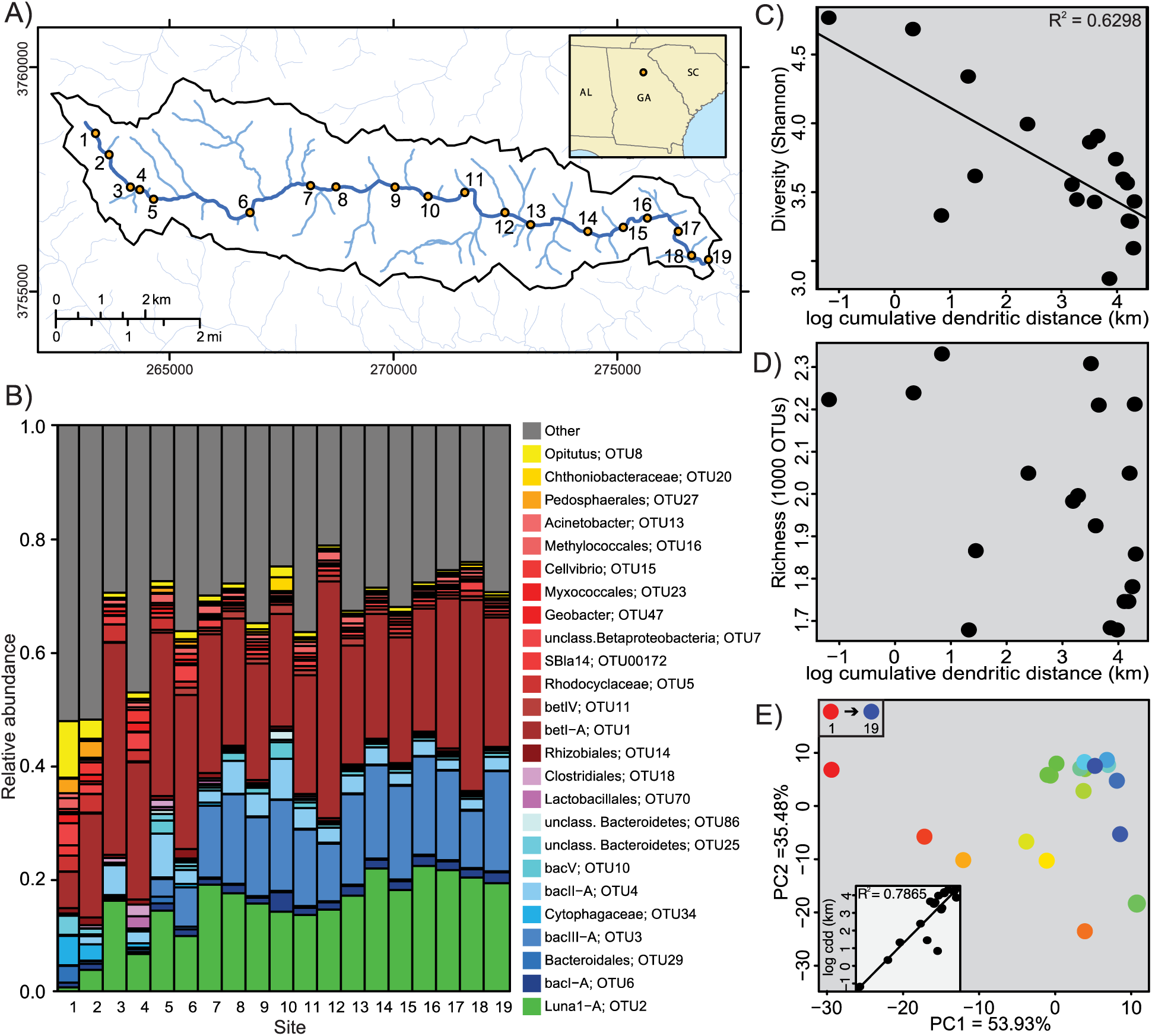
Transitions in bacterial communities from the headwaters of McNutt Creek to downstream sites. A) The flow path of McNutt creek is shown in dark blue and all samples were taken along its length, noted by the numbered collection sites. Tributaries are shown in light blue. B) The relative abundance of OTUs at each location. All OTUs representing 1.5% or more of the community in any sample in the full study (including samples not shown here) are displayed. All taxa falling beneath this threshold were group into the other category in grey. Samples are aligned by their location along the creek, starting at the headwaters, as shown in A. Shannon diversity (C) and OTU richness (D) is shown for each sample and displayed according to the cumulative dendritic distance at that location in the stream. E) Ordination of samples along PC1 and 2 are display in color according to their location along the stream path, with upstream-most samples in red, working down the color spectrum to purple. Inset displays PC1 values for each site plotted against cumulative dendritic distance.

## Methods

### Sampling schemes

Pelagic water samples were collected from McNutt Creek, Athens, GA, USA (33°55’50.29“N, 83°30’30.91“W). Nineteen sampling locations were selected based on their location in the stream network (headwaters versus main stem) and the ability to access these sites, as some were located on private property and required the consent of the owners. Water samples were collected in 2016 and 2017 to assess community assembly and stability along the headwater (1^st^ order) to main stem (3^rd^ order) longitudinal gradient. In 2016, three streams (sites 3, 12, and 19) were sampled daily from June 9^th^ to June 19^th^. On June 14^th^, all nineteen sites were sampled within 7 hours. Three groups of three to four volunteers each were sent out in groups to survey a third of the creek, working from the most upstream site downstream.

At each site, water was collected mid-stream from the water column approximately at mid-depth in 4 L acid-washed cubitainers. Each container was rinsed with stream water three times before the final sample was collected. Water was processed either immediately on-site or within an hour of collection following the sampling methods outlined in Hassell et al. (10). Samples were run through sterilized tubing with in-line filtration through a 5.0 μm, 47 mm diameter SVPP pre-filter (Millipore) to capture particulate matter. The effluent was then run through a 0.22um, 2 ml sterivex filter (Millipore). A total of 500ml of water was filtered for each sample. Filters were preserved at -80 °C until DNA extraction. In addition to the filtered water samples, a YSI Professional Plus meter was used to temperature, pH, dissolved oxygen percentage, and conductivity (Table S2). Dissolved organic carbon (DOC), total dissolved nitrogen (TDN), and total dissolved phosphorus (TDP) were measured for selected filtered water samples by the Stable Isotope Ecology Lab at the University of Georgia (Table S2). Daily air temperature measurements were available through Weather History for Athens (https://www.wunderground.com/history/airport/KAHN/) and daily precipitation measures were available through the National Weather Service (https://www.weather.gov/ffc/rainfallscorecard).

In 2017, sampling was focused on headwater-proximal sites (Sites 1-6) to determine the reproducibility of initial community assembly. Sites 1, 3, and 6 were sampled daily from August 24^th^ to September 3^rd^. On August 24^th^, August 29^th^, and September 3^rd^ samples were collected from all six upper-most sites (sites 1-6). The 2017 samples were collected using the same methods as used in 2016.

### Watershed Characteristics

Geographic information system (GIS) analysis was used to determine physical differences between each site’s watershed, drainage network, and land use/land cover. ArcMap 10.5 was used to calculate each sample site’s watershed area, total cumulative dendritic distance (CDD), land use/land cover characteristics, and impervious surface area. Each sample point’s watershed was delineated using “Hydrology” tools in the Spatial Analyst toolbox and the National Elevation Dataset’s 30 m digital elevation model (DEM) (26). Flow direction and flow accumulation rasters were created from a hydrologically-corrected DEM (“Fill” tool was used to correct the DEM). After the sampling points were associated with the flow accumulation raster (using the ‘Snap Pour Point’ tool), each watershed was delineated using the ‘Watershed’ tool. Total dendritic distance was calculated by using each sampling point’s watershed to clip the high-resolution National Hydrography Dataset’s stream layer (27). Similarly, each watershed was used to extract land use/land cover characteristics and impervious surface cover from the 2011 National Land Cover Database (28, 29).

### DNA extraction

DNA was extracted from the sterivex filters following the methods outlined in Hassell et al. (2018) (10). Filters were thawed, then 1ml of lysis buffer (40 mM EDTA, 50 mM Tris (pH 8.3), 0.73 M Sucrose) and lysosome dissolved in lysis buffer (2.11 mg/ml final concentration) were added and incubated at 37° C for 30 min while rotating. Proteinase K dissolved in lysis buffer (0.79 mg/ml final concentration) and 200ul 10% SDS was added for a second incubation step at 55° C for 2 h. Lysate was extracted and mixed with an equal volume of Phenol:Chloroform:IAA (25:24:1; pH 8.0). Samples were centrifuged for 5 min at 3500xG and the top phase was saved. 0.04 x volume 5M NaCl and 0.7 x volume isopropanol was added, mixed, and incubated at room temperature for 10 min. Samples were centrifuged for 15 min at 17,000xG and supernatant was discarded. DNA was resuspended in 400ul elution buffer (Omega Biotek, Norcross, GA, USA) and incubated at 65° C while rotating for 10 min. DNA was then processed using Omega Biotek’s E.Z.N.A. Water DNA kit following manufacturer protocol from step 13 through completion (Omega Biotek, May 2013 version).

### Sequencing and analysis

Following the methods outlined in Tinker and Ottesen (30), the V4 region of the 16S rRNA gene was amplified in each DNA sample. The resulting library was submitted to the Georgia Genomics Facility for sequencing (Illumina MiSeq 250 × 250 bp; Illumina, Inc., San Diego, CA). The resulting raw sequences from these experiments are available under the accession numbers SRP155540 in the NCBI Sequence Read Archive.

Returned reads were processed using the mothur software package (31) following the MiSeq standard operating protocol with the following modifications: reads that fell outside of 200-275bp were excluded from contig generation; SILVA reference database release v123 was used for sequence alignment; primers GTGCCAGCMGCC-GCGGTAA and GGACTACHVGGGTWTCTAAT to perform in-silico PCR; the VSEARCH algorithm was used to identify chimeric sequences (32); taxonomic classification was completed via the Wang Method using the May 5^th^, 2013 release of the greengenes reference database, version 13_8, in combination with the FreshTrain dataset following the TaxAss workflow (33-35); during taxonomic assignment a bootstrap value of 70 was used; an additional remove.lineage command was run to ensure that sequences from cyanobacteria, chloroplasts, and unknown taxa were removed. Operational taxonomic units (OTUs) were called at 97% or greater sequence similarity. From the 91 samples that were collected from these studies, a total of 6 119 579 of 16S rRNA gene sequences passed quality filtering steps, resulting in an average of 67 248 sequence reads and 1 562 OTUs per sample (Table S1).

Statistical analysis was completed in R using the vegan package (36). When necessary for analysis, samples were rarefied to a depth of 3 291 sequences. The envfit function was used to calculate significant (p ≤ 0.01) physiochemical parameters and watershed characteristics that are displayed as loadings on ordination plots. The cor.test function was utilized to test for significance of Spearman’s correlations. T-tests were used to determine significance between boxplots.

## Results

### Study site and physicochemical data

McNutt Creek is a 20 km-long stream in Athens, GA, USA that flows through a mixed land use area, spanning agricultural, residential and forested sections (Fig. 1A, Table S2). Stream width ranges from 1.10 m (site 1) to 5.18 m (site 19). McNutt Creek is part of the Upper Oconee Watershed, a temperate, urban watershed that provides drinking water for the city of Athens and the surrounding area. Two studies of temporal and spatial dynamics in microbial community composition in McNutt Creek were conducted.

The first study (in 2016) was an 11-day daily collection of samples at 19 sites along the entire creek length. Water was also collected at three sites (sites 3, 12, 18) for the five days immediately preceding and following the 19-site collection to survey daily community shifts along the creek length. During the 2016 study period, 9 June through June 19, the average daily high and low air temperatures were 33.69 °C and 20.20 °C, respectively (Table S2). The average water temperature (22.96 °C) generally increased downstream, with a minimum of 20.15 °C (site 2) and a maximum of 24.98 °C (site 18). There was measurable precipitation on June 11^th^ (0.33 cm), 12^th^ (0.31 cm), 14^th^ (1.56 cm), and 17^th^ (0.20 cm) for a total of 2.39 cm of rain during the 2016 study. All samples collected on June 14 were taken before the rainfall, with the exception of site 4.

TDP, TDN, and DOC were measured for samples collected on June 14^th^. TDP ranged from 11.28-49.24 mg/L with an average of 24.84 mg/L, and no trend was observed along the flow path. TDN fluctuated among the headwater-proximal sites before stabilizing around 0.6 mg/L. Average TDN was 0.64 mg/L, with a range of 0.45-1.02 mg/L. DOC was relative stable at all sites, with the exception of site 7, and averaged at 1.45 mg/L, with a minimum of 0.99 at site 16 and 6.44 mg/L at site 7.

To address fluctuations in community profiles at the headwaters, a second study in 2017 focused on the six most upstream sites (sites 1-6). Water was collected from each upstream site on study days 1, 6, and 11. To understand daily changes in headwater community composition, sites 1, 3, and 6 were sampled daily for 11 days. Average daily air temperatures were slightly lower than observed during the 2016 study, with an average high and low of 29.14 and 18.73 °C, respectively (Table S2) (37). Average water temperature was 20.15 °C, with minimum and maximum water temperatures of 18.53 (site 1) and 21.59 °C (site 6). A total of 3.03 cm of rainfall fell during the 2017 study, with measurable precipitation occurring on August 30^th^ (0.43 cm) and 31^st^ (2.59 cm). TDN, TDP, and DOC were measured at site 3 on all days, and averaged 0.48, 29.51, and 1.33 mg/L respectively, comparable to measurements taken in 2016.

### Longitudinal study of McNutt Creek

In the 2016 longitudinal study, alpha diversity (Shannon’s) significantly decreased with increasing cumulative dendritic distance (CDD; p < 1×10^-4^, rho = 0.63) (Fig. 1C). The microbial community at the headwaters (site 1) was highly diverse, with 47.44% of sequences belonging to taxa present at < 1.5% of the community (Fig. 1B). The fraction of the population defined by lower-abundance taxa (“others”, <1.5%) progressively decreased with increasing CDD, shifting from 52.26% at site 1 to 29.50% at site 19, with a minimum of 21.28% sequences present at site 12.

To explore changes in microbial community structure and beta diversity, principal components analysis (PCA) was completed using rarified samples (Fig. 1E). Log-transformed total CDD correlates strongly with the position of samples along the first principal component (Fig 1E inset, p < 1×10^-4^, R^2^ = 0.79). Upstream samples are scattered but downstream sites are clustered more tightly, suggesting a decrease in variance with increasing CDD. Incorporation of physiochemical, land use, and nutrient data suggest that tree cover type (deciduous upstream, evergreen downstream), dissolved oxygen levels, and urbanization factors may influence microbial community structure (Fig. S1). However, given that only a single linked flow path is being examined, these relationships are difficult to interpret and may be autocorrelative.

To directly assess distance-decay relationships along the length of the stream, Bray Curtis dissimilarity (weighted and unweighted) was calculated between each sample site and plotted against the length of stream between them (Fig. S2). These relationships were calculated in terms of upstream and downstream proximity. For both weighted and unweighted measures, 12 relationships were significant (p < 0.05). All sites downstream of site 13 displayed significant dissimilarity between adjacent sites, indicating a downstream increase in similarity decay.

In an analysis of the 250 most abundant taxa, a statistically significant negative correlation (Spearman, p < 0.05) was found between the relative abundances of 76 OTUs and CDD (Fig 2). Together, negatively correlated taxa represent 49.16% of sequences in headwater site and 6.58% of sequences at site 19 (Fig. 2B). In contrast, a statistically significant positive correlation was found between 36 OTUs and total CDD (Fig. 2A). Positively correlated taxa include OTUs belonging to the freshwater-associated Actinobacteria clade Luna1-A OTU2, and Bacteroidetes clades bacIII-A OTU3 and bacI-A OTU6 (nomenclature from Newton et al., (38)) (Fig. 2A). All positively correlated taxa together represent 4.44% of sequences in the furthest headwater site (Site 1) and 45.81% of sequences in the furthest downstream site (Site 19).

**Figure 2.**
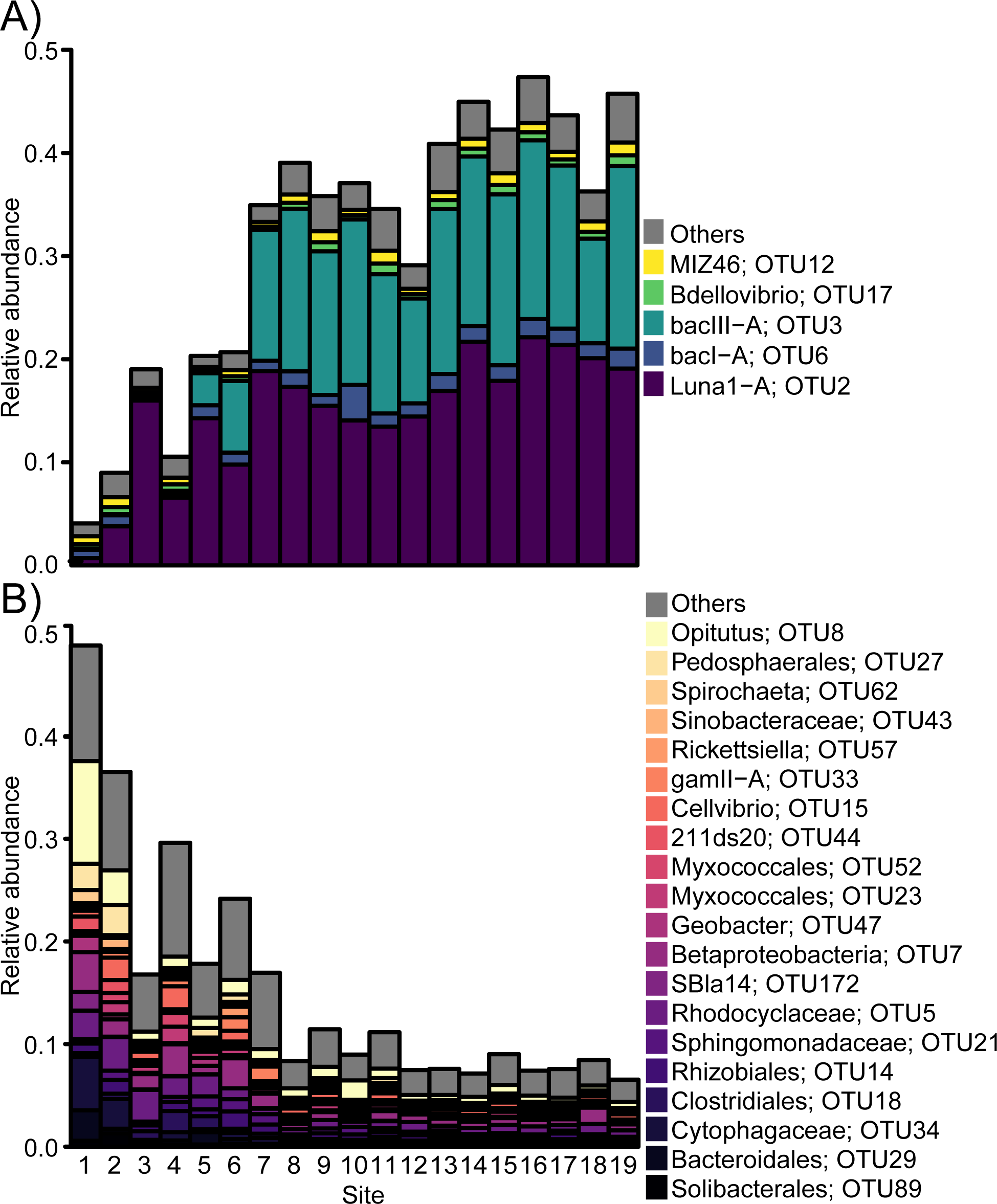
Changes in the positively and negatively correlatied OTUs along the stream length. The relative abundance of all significantly positively (A) or negatively (B) correlated taxa comprising at least 1% of any sample are displayed in color. Alladditional taxa below this threshold are reported as “others” in grey. OTU numbers are reported to the right of all taxa for reference.

### Daily fluctuations in community composition at the upper, middle, and lower reaches of McNutt Creek

Daily assessment of sites 3, 12, and 18 during June 2016 revealed that the taxonomic profile of the community at each site was relatively stable at the OTU level (Fig. 3A). For site 3, 675 OTUs (out of 14 860 observed) were shared across the entire 11-day time series, representing an average of 96.11% (range 95.12 -97.20%) of recovered sequences. For site 12, 513 OTUs were shared across the sampling period (out of 15 022), representing an average of 96.75% (range 95.47 -97.64%) of recovered sequences. For site 18, 413 OTUs were shared across the 11-day time series (out of 15 122), representing an average of 95.17% (range 88.30-97.47%) of recovered sequences. Across all dates and times, 344 OTUs were shared, representing 93.61% of recovered sequences.

**Figure 3:**
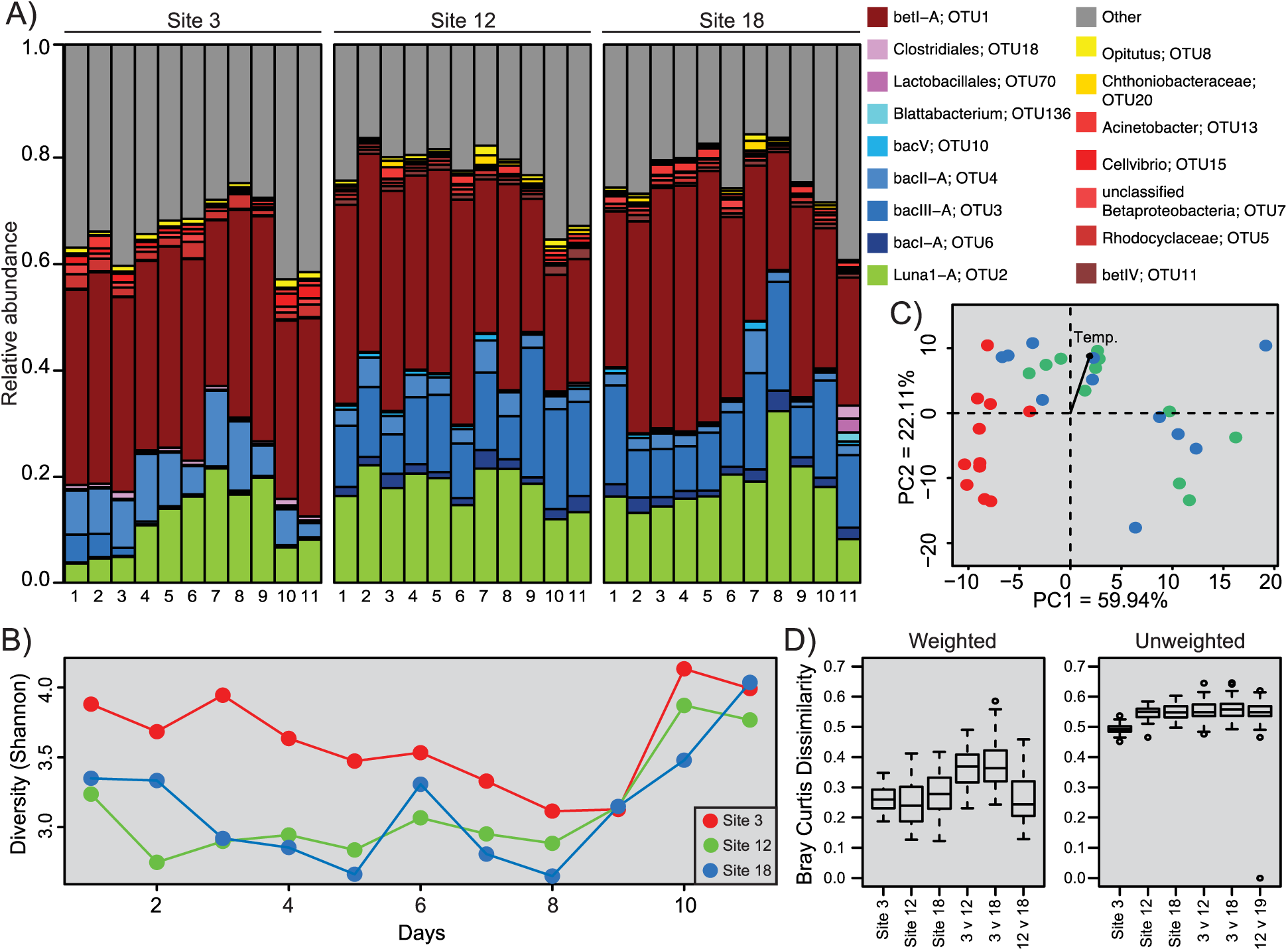
Daily analysis of bacterioplankton communities across the length of the creek. A) The relative abundance of OTUs present at >1.5% of any one sample is displayed (all other OTUs are grouped under ’other’). B) Shannon diversity plotted against day of sampling for each of the three selected sites, with site 1, 12 and 18 shown by red, green and blue dots, respectively. This color scheme was used to show sample ordination in the PCA plot displayed in C), in which envfit was used to calculate loadings of the parameters listed in Table S1. D) Bray Curtis dissimilarity was determined for and between all samples taken from each site.

Alpha diversity measurements fluctuated at all sites throughout the study (Fig. 3B). The bacterial community at site 3 was generally the most diverse, showing the highest Shannon index value of the three sites in 9 of 11 days. Site 3’s Shannon diversity was significantly higher than both other sites, with p-values of 0.004 (site 12) and 0.008 (site 18). Differences in Shannon index between sites 12 and 18 were smaller and not statistically significant. The relationship between these sites were also less predictable, with 18 exhibiting a Shannon Index less than 12 for only 4 of 11 days.

Beta diversity analyses suggest that site 3 hosted a distinct community from downstream sites. Weighted Bray-Curtis analyses show the median dissimilarities between site 3 and the two downstream sites are markedly higher than the median dissimilarity between sites 12 and 18. Similarly, PCA analysis shows a clear separation of site 3 from the downstream samples (Fig. 3C), while samples from sites 12 and 18 cluster together. Though there were no significant differences between site 3’s ordination points and site 12’s or 18’s ordination points, significant differences in water temperature may be driving this separation (p < 0.001). Analysis of Bray Curtis dissimilarities between successive samplings at the same site shows temporal trends in community similarity over time for abundance-weighted (p = 0.012 [site 3], 0.013 [site 18]) but not unweighted (presence/absence) measures (Fig. S3). No significance was found for either measure at site 12.

To understand which taxa consistently increase or decrease downstream, downstream trends in the abundance of the 250 most-abundant taxa were analyzed. 213 were found to decrease downstream on at least one day. ZB2 OTU208, ABY1 OTU280, unclassified Rhodospirillaceae OTU38, and bacII-A OTU4 all decreased downstream on 10 of the 11 sampling days. In contrast, 94 of the 250 were identified as increasing on at least one day, although only 12 increased on at least 5 days. BacI-A OTU6 and bacIII-A OTU3 were found to increase between sites 3, 12, and 18 during multiple days. Alphaproteobacterium Alf-V OTU193 was the most consistent OTU of the top 250, increasing down the stream length for 9 of the sampling days. Other organisms, such as Luna1-A OTU2, bacV OTU10, betI-A OTU1, and betIV OTU11, that increased along the full flow path in the 19-site study only exhibited increases across these three sites for a few of the individual sampling days.

### Temporal and spatial trends in community assembly at headwater proximal sites

The 2017 spatial and temporal study was aimed at examining community assembly near the stream headwaters in greater temporal and spatial detail. Three longitudinal studies of the 6 most uppermost stream sites were performed at 5-day intervals to evaluate longitudinal trends in community composition (Fig. 4). Across these three collections, qualitatively similar trends in taxonomic composition were observed (Fig. 4A). Alpha diversity was significantly correlated with log CDD on the second (but not the first or third) longitudinal sample collection (p = 0.0045) (Fig. 4B). Paralleling the results from the full stream length study in 2016, more taxa significantly decreased (93 OTUs showed negative relationships on at least 1 of the three days) along the flow path than increase (9 OTUs), but those that increase in abundance comprised a substantial fraction of the total population by sites 5 and 6 (Fig. S4). 35 of the 93 taxa exhibiting significant negative relationships and two of the nine taxa exhibiting positive relationships (Luna1-A OTU2 and bacIII-A OTU3) with CDD also did so in the 2016 study (Fig. S4, Fig. 2).

**Figure 4.**
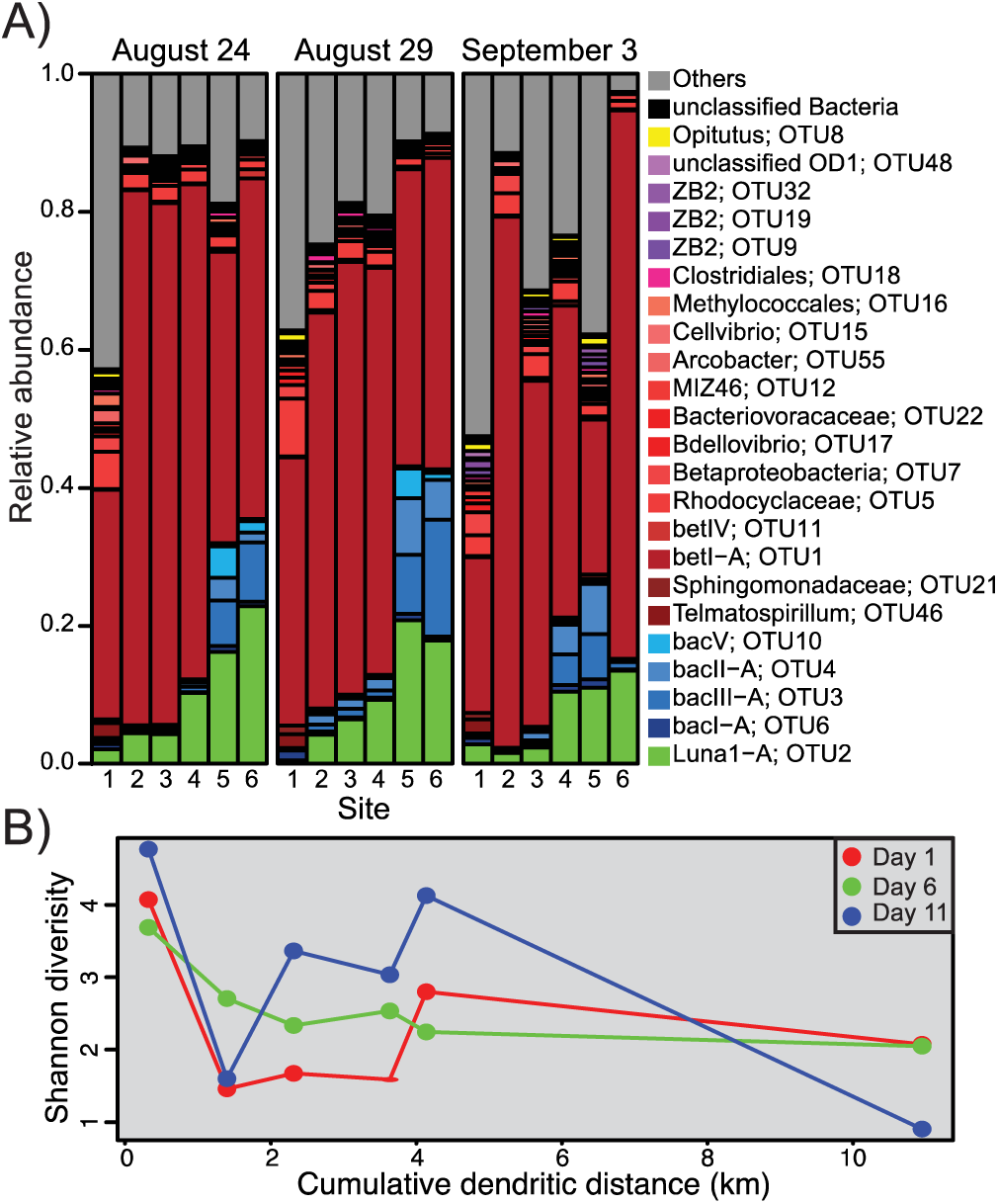
Community analysis of all headwater site samples collected between at three time points in 2017. A) The relative abundance of taxa present in each sample is displyaed according to the parameters defined in Fig. 1. B) Diveristy for each sample plotted according to the cumulaive dendritic distance measurement for each site.

Daily fluctuations in microbial community composition were evaluated at sites 1, 3, and 6 from August 24-September 3, 2017 (Fig. 5A). Alpha and beta diversity were consistently greater at site 1 and lower at sites 3 and 6, in a manner similar to that of the whole stream (Fig. 1C). Site 1 was found to be significantly more diverse (Shannon, p = 0.0057, 1.077×10^-5^), than sites 3 and 6, which were not significantly different. Day-to-day beta dissimilarity was assessed for each focus site (site 1, 3, and 6) to understand community change over time (Fig. 5D). Median weighted Bray Curtis dissimilarity across time was higher at site 1 than either downstream site, significantly different from site 3 by presence-absence measures (p = 0.0219) and from site 6 by both measures (p = 0.0034, <1×10^-4^). Median unweighted dissimilarity was greatest and most variable at site 6. At site 1, 197 OTUs out of 15 338 were shared across the entire 11-day time series, representing an average of 87.11% (range 81.83- 94.35%) of sequences recovered from this site. For site 3, 97 (of 15 438) were shared across the sampling period, representing an average of 86.31% (range of 71.42-98.70%). This was substantially lower than the 675 OTUs shared at this site during the 2016 daily study. For Site 6, 34 OTUs (of 15 501) were shared across the 11-day time series, representing an average of 91.06% (range 76.18-98.65%) of recovered sequences.

**Figure 5.**
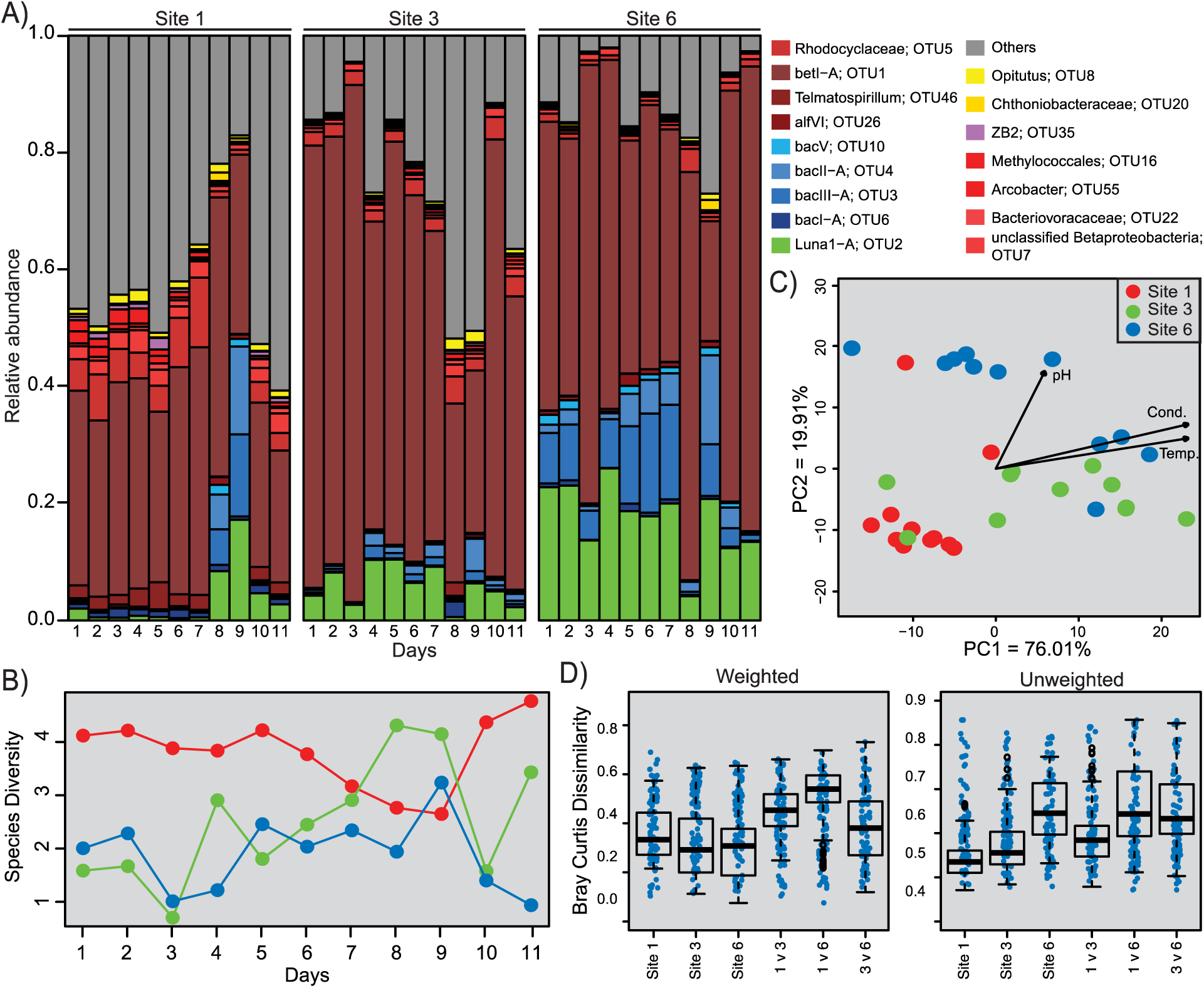
Analysis of daily water samples collected from three headwaters sites across multiple days. A) Relative abundance of all taxa present in each sample is displayed according to parameters defined in Fig. 1. B) Community diversity observed at each site over the course of the study, with samples from site 1, 3 and 6 represented by red, green and blue points, respectively. C) Ordination of all daily samples in the resulting PCA plot. D) Bray Curtis dissimilarity was determined for and between all samples taken from each site. Blue dots represent individual samples and open circles represent outliers

In a PCA of these data (Fig. 5C), samples clustered by site, suggesting each location had a distinct microbial community, potentially driven by significant physiochemical differences (pH, p < 1×10^-4^; conductivity, p = 0.0006; temperature, p = 0.0006). PERMANOVA found a significant effect of site but not sampling day on community composition (p = 0.001 for site, 0.12 for day). No individual sites exhibited significant temporal distance-decay relationships in Bray-Curtis dissimilarity, with samples taken 24 hours apart showing similar levels of dissimilarity as to samples taken up to 5 days apart (Fig. S5).

All three sites were consistently dominated by bet1-A OTU1 during the study period, and no consistent trend in its abundance was observed between sites. However, other Proteobacteria showed positional trends in representation. For example, alfVI OTU26 and betIV OTU11, were found to increase on 5 days, though they showed decreases on a single study day. Other proteobacterial OTUs were noted to decrease over the study period, including Telamatospirillum OTU46, Bdellovibrio OTU17, and Geobacter OTU47. As was observed in 2016, Luna1-A OTU2, and bacIII-A OTU3 increased across the three sites on 8 or more of the study days. Interestingly, bacII-A OTU4 exhibited a positive trend with total CDD on 6 of the 11 sampling days in 2017. These results are in stark contrast to the negative trends this OTU exhibited for 10 out of 11 days in 2016.

The 2.54 cm rain event captured between day 7 and 8 (August 30^th^ and 31^st^) had variable effects on the abundance of prominent taxa at each site. As mentioned above, several Proteobacteria, which increased along the sampled flow path on other days, decreased on either day 8 or 9. This was also the case for bacV OTU10, though this OTU had inconsistencies in trend direction in the 2016 study. Taxa that typically showed positive trends with CDD (e.g. Luna1-A OTU2, bacIII-A OTU3, bacII-A OTU4), did not on days 8 or 9 but returned to exhibiting trends in increasing on day 10. Days 8 and 9 also showed differences in the relative prevalence of low-abundance taxa (“others”) at each site: site 3, for example, averaged 51.14% on days 8 and 9 and 19% across the rest of the time series.

### Comparisons of community composition at daily to annual time scales

In the August 2017 headwater time series, Proteobacteria were dominant across all days and sites at substantially higher abundances than observed in the 2016 study (Fig. 4, 5). Proteobacteria represented 54.56% of sequences at site 3 in 2016 and 67.33% in 2017 during both 11-day sampling periods. Bacteroidetes and Actinobacteria, in contrast, were less abundant (average of 13.80% for Bacteroidetes and 11.55 % for Actinobacteria in 2016 vs. 6.88% and 6.12%, respectively, in 2017). However, as in 2016, these phyla were observed to consistently rise in abundance with increased CDD (Fig. 4 and Fig. 5). At the OTU level, overlap across years was relatively low, with only 25 of the 8 943 OTUs detected across all samples in 2017 also present in all samples collected in 2016. However, these shared OTUs were among the most abundant sequences identified, including 24 of the 100 most abundant OTUs.

Because daily samples were collected at site 3 in both 2016 and 2017, we were able to directly compare community similarity across both daily and annual time scales (Fig. 6). The 2017 daily time series exhibited significantly greater day-to-day dissimilarity by both weighted and unweighted measures than observed in 2016 (t-test, p = 0.01 and p < 1×10^-4^, respectively). In fact, day-to-day variability in 2017 was similar in scale to cross-year variability between 2016 and 2017. To account for the potential effects of the rain event on day 8 of the 2017 study, significance was re-evaluated by removing days 8 and 9 from the data set and rerunning dissimilarity calculations. Weighted dissimilarity was still significantly different although less so (p = 0.03). Unweighted dissimilarity, however, was no longer significant (p = 0.97). Principal components analysis shows some degree of clustering within years, with the 2017 samples largely separating along the first principal component (87.58% of variation), and 2016 samples separating along the second principal component (9.05% of variation), although no metadata besides collection year (envfit, p = 2×10^-4^) was found to be significantly different between the two samples sets.

**Figure 6.**
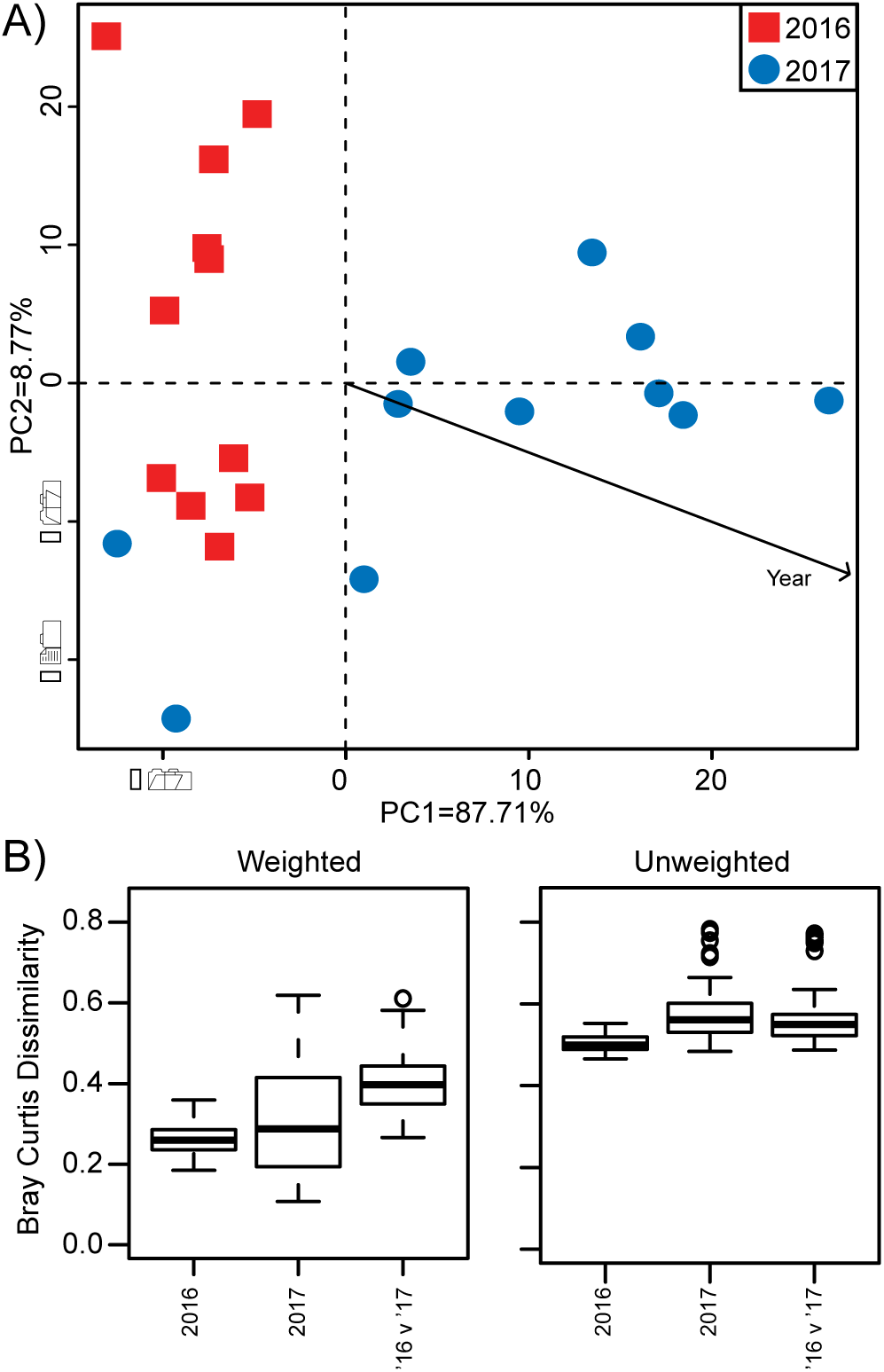
Comparison of site 3 across years. A) Resulting ordination of PCA analysis. Envfit was run with daily metadata but no values were significant. B) Bray Curtis dissimilarity was calculated for samples from each year and between years.

## Discussion

This work examined the bacterial community assembly at a daily resolution in a third-order stream located in a temperate, urban watershed. While pelagic freshwater bacterioplankton communities have been previously described around the world, the stability and renewal of these populations in headwater streams remains in question. In particular, we aimed to elucidate, on fine temporal and spatial scales, the degree to which lower-order streams mimicked community assemblages and diversity trends observed in higher order rivers and whole watersheds (1, 2, 4, 8, 9, 11).

Along the length of McNutt Creek, an inverse relationship between alpha diversity and CDD was observed, which has been previously reported in other river systems (9, 10, 12, 23). Patterns of decreasing alpha diversity with downstream distance traveled were most apparent across the full stream length (Spearman’s R^2^ = 0.6298 p = 0.005), however, site-to-site fluctuations were apparent in both daily studies. It is also of note that alpha diversity calculations for the 2017 study were typically lower than those recorded in 2016, although the cause of this diversity loss is unknown.

Downstream trends in site-to-site beta diversity comparisons were present but relatively weak, primarily exhibiting differences between the headwater and furthest sites downstream. For example, significant beta diversity relationships were found between site 3 and 18, and between sites 12 and 18 in the 2016 study. When dissimilarity was assessed along the entire stream length on a site-by-site basis, the majority of sites were found to have a significant positive correlation with distance, particularly at the extreme ends of the stream (Fig. S3). During 2017, the only significant beta diversity relationships were between site 1 and both downstream sites, 3 and 6. These results confirm earlier findings in the broader Upper Oconee watershed, where three of five seasons exhibited trends in beta diversity loss with increasing cumulative dendritic distance (10). This pattern of dissimilarity loss with increasing hydrologic distance is not unique to the Upper Oconee watershed, and was also observed across the Ybbs river network in Austria (2).

The effects of time between samples collected on community structure were apparent at some but not all sites, suggesting a limited impact of the previous community composition on later ones. A similar trend in diversity was recorded in rock biofilms in a headwater stream, in which weekly samples from the same site were less similar than those taken from other locations along the creek, although site locations were considerably closer together than in the present work (39). As dispersal in streams is primarily unilateral as microbes are passively transported, this speaks to the strength of renewal of these communities and their resilience to immigrant taxa from neighboring environments.

Water temperature is the sole environmental factor that was found to have a statistically significant correlation with community composition over both time and space, suggesting that water temperature may play a key role in shaping stream community composition. None of the other physiochemical parameters measured in this study showed consistent, statistically significant correlations with community progression or composition. These results are not entirely surprising, given the results of a previous study on the greater Upper Oconee watershed, which found that position within the watershed showed far more significant impacts on community composition than individual physicochemical factors (10). In addition, studies in other fluvial and freshwater systems have found substantial temporal and spatial variability in the relative importance of different physiochemical parameters in shaping community composition (25, 40). Land use parameters associated with development were significantly associated with community composition across sites (Fig. S1), and similar relationships were also noted in de Olivera and Margis (13). However, in this study, these relationships are more likely to be a coincidental finding due to the fact that downstream portions of this stream happened to be more highly developed.

During the 2017 sampling period of headwater sites, a storm event resulting in 2.54 cm of rainfall occurred. Rainfall has previously been shown to introduce taxa into freshwater systems and increase diversity (22, 41, 42). While our results are consistent with this, it is interesting that each of the three sites sampled appeared to be affected differently by the influx of rainwater. At site 1, we observed a notable loss of alpha diversity paired with the introduction of Bacteroidetes and Actinobacteria, while site 3 exhibited a noticeable increase in total diversity, and site 6 exhibited a decrease in diversity and an increase in the fraction of Proteobacteria (Fig 2). All sites returned to post-rain community compositions within 48 hours (Fig. 4). Several other minor precipitation events occurred during both studies however no strong disturbance was detected in the community data. While further studies are merited to better understand both the consistency in disruption and recovery of bacterioplankton populations along the entire stream reach, these data suggest that headwater bacterial communities are highly resilient to rainwater influx.

Throughout both 2016 and 2017 studies, a small set of taxa found at low abundance in the headwaters of the stream was enriched in downstream sites. This enrichment corresponded with the significant decrease of many OTUs prevalent at the headwater sites. These findings support the hypothesis that freshwater streams function as major site of selection and species sorting (2, 8, 10, 12). The set of taxa that were specifically enriched in downstream environments was relatively consistent over time. Multiple OTUs were identified as significantly positively correlated with stream length across days and years, 36 and 9 of the 250 most abundant taxa in 2016 and 2017 respectively. Many of these OTUs belonged to well-known ‘typical’ freshwater bacterial clades (10, 23, 25, 38), including: Luna1-A (OTU2), bacIII-A (OTU3), bacVI (OTU90), bacV (OTU10), betI-A (OTU1), betIV (OTU11), and bacI-A (OTU6). It is of note that in the 2016 study, a shift in prevalent Bacteroidetes members from bacII-A OTU4 in upstream environments to bacI-A OTU6 and bacIII-A OTU3 further downstream occurred, suggesting that certain Bacteroidetes clades are better adapted either for life in different regions of the stream or for long-term exposure in freshwater environments. Interestingly, bacII-A OTU4 was identified as significantly positively associated with CDD in 2017, when only the upper reaches of the stream were examined.

These data parallel findings in the existing literature in which a consistent freshwater stream microbiome has been proposed (2, 9, 10, 13, 19, 21, 43), though the exact definition of this population and the nomenclature describing this phenomenon has been highly variable. For example, the “core” freshwater community of stream biofilms and sediment was defined by Besemer et al (2013) as taxa found in at least 50% of all samples (2). A stricter definition was employed in de Olivera and Margis’ seasonal study of an entire river length, describing the “core” community as taxa that persisted across all samples and timepoints (13). In contrast, Ruiz-Gonzalez et al. (21) defined the ‘core seed bank’ of bacterial taxa as those that were more abundant in downstream than upstream sites vs. ‘restricted’ taxa that decreased in abundance with increasing downstream distance. (21). It is clear that a select and consistent set of taxa dominate freshwater habitats in fluvial systems at all scales (23, 25, 40, 43). However, it is also clear that the abundances of these microbes can vary substantially, both within and between fluvial networks (40). While these variations are often posited to be driven by physicochemical variables, the relationships found between specific taxa and different environmental variables have often varied across studies (4, 16), and in some cases even across time within a single system (19). With these and other works, it becomes increasingly apparent that the concept of a ‘typical’ freshwater microbiome (25) extends across all levels of the freshwater continuum, with even a single stream exhibiting strong and reproducible selection for a small subset of microbial taxa that exhibit variable abundances over time.

## Conclusions

In this work, we observed the rapid and consistent assembly of freshwater bacterioplankton communities in a small, temperate headwater creek, paralleling population trends reported in the entire watershed and in systems around the globe. Downstream travel was associated with a consistent enrichment of a set of bacterial taxa belonging to clades that are widely associated with freshwater environments (38). As these clades rise to dominate the bacterioplankton population, a larger number of taxa prevalent at the head of the stream decrease in abundance. These shifts in community composition gave rise to consistent trends in overall alpha and beta diversity along the length of the stream. However, substantial variability in community composition among low-abundance taxa was maintained at each site, with changes in community composition over 24 hours that in some cases mirrored dissimilarity among samples collected up to 1 year apart. Overall, these results underpin the importance of headwaters as the site of rapid in-stream selection that gives rise to a highly consistent, site-specific stream microbiome that dominates the water column of downstream environments.

